# White matter changes in the perforant path in patients with amyotrophic lateral sclerosis

**DOI:** 10.1101/547588

**Authors:** J. Mollink, M. Hiemstra, K.L. Miller, I.N. Huszar, M. Jenkinson, J. Raaphorst, M. Wiesmann, O. Ansorge, M. Pallebage-Gamarallage, A.M. van Cappellen van Walsum

**Author notes:** Equal contribution (J. Mollink, M. Hiemstra).

## Abstract

Amyotrophic lateral sclerosis (ALS) is a progressive and incurable motor neuron disease. Some ALS patients are affected by a level of cognitive or behavioural decline that meets the criteria for frontotemporal dementia (FTD). ALS and FTD share genetic and pathological features; for example, the deposition of phosphorylated 43 kDa TAR DNA-binding protein (pTDP-43) in the brain. Spreading of pTDP-43 pathology in ALS towards brain areas that connect via the Papez circuit is a possible indicator of progression towards FTD. For example, pTDP-43 aggregates in the granule cells of the hippocampus correlate well with clinically manifest FTD. Here, we test the hypothesis that white matter degeneration of the perforant path – as part of the Papez circuit – in the hippocampus is a feature of ALS, even in the absence of fully developed FTD or deposition of pTDP-43 inclusions in hippocampal granule cells. We used diffusion MRI (dMRI), polarized light imaging (PLI) and immunohistochemical analysis of hippocampus sections from controls (n=5) and ALS patients (n=14) to perform an in-depth study of white matter in the perforant path.

The dMRI results show a significant decrease in fractional anisotropy (p=0.01) and an increase in mean diffusivity (p=0.01), axial diffusivity (p=0.03) and radial diffusivity (p=0.03) in the perforant path in ALS patients compared to controls, possibly indicating a loss of white matter fibres. Myelin density (measured with PLI retardance) was lower in ALS patients compared to controls (p=0.05) and correlated with dMRI fractional anisotropy (r=0.52, p=0.03). The dMRI and PLI results were confirmed by the immunohistochemistry; both myelin (proteolipid protein, p=0.03) and neurofilaments (SMI-312, p=0.02) were lower in ALS patients. The activated microglial (CD68) density was similar in ALS and controls. Only two out of the fourteen ALS cases showed pTDP-43 pathology in the dentate gyrus; however, while these two ALS-FTD cases showed reduced myelination in the perforant path, the values were comparable to other ALS cases.

We conclude that degeneration of the perforant path occurs in ALS patients and that this may occur before, or independent of, pTDP-43 aggregation in the dentate gyrus of the hippocampus. Future research should focus on correlating the degree of clinically observed cognitive decline to the amount of white matter atrophy in the perforant path.

## Introduction

Amyotrophic lateral sclerosis (ALS) is a motor neuron disease that is primarily characterized by a loss of motor neurons in the brain and spinal cord causing dysfunction of voluntary movement. A significant proportion of ALS patients develop cognitive impairment in the form of frontotemporal dementia (FTD). It is estimated that up to ∼13% of the patients with ALS meet the criteria for FTD, with an additional ∼35% demonstrating some form of cognitive impairment [43, 47]. In addition to the symptomatic overlap between ALS and FTD, pathological aggregation of phosphorylated 43 kDa TAR DNA-binding protein (pTDP-43) [2, 45] is reported in almost all cases of ALS and in the largest subset of FTD (behavioural variant FTD: bvFTD) supporting the concept that ALS and FTD are part of the same pathological spectrum [15, 45].

A pathological staging system in ALS and bvFTD, based on the regional distribution of pTDP-43, highlights the involvement of several cortical regions that connect with each other via the Papez circuit in both diseases [15, 16, 61]. The Papez circuit (Fig. 1) – comprised of hippocampal formation, mammillary bodies, anterior thalamic nuclei and anterior cingulate – is a principle pathway of the limbic system associated with the formation and consolidation of episodic memory [46, 58]. Recent evidence recognises episodic memory deficits in bvFTD [29]. These deficits are supported by in-vivo MRI volumetric analyses that demonstrate regional grey and white matter atrophy of the Papez circuit [30, 32]. Furthermore, volumetric analysis on post-mortem brains demonstrated reduced grey matter volume, while white matter connectivity was not investigated [30]. Less is known about the involvement of the Papez circuitry in patients with ALS. Previous studies have demonstrated a relationship between pTDP-43 pathology in ALS patients and their cognitive decline (Brettschneider *et al*., 2013; Cykowski *et al*., 2017), suggesting a relation to limbic system impairment. Furthermore, an accumulating body of evidence supports involvement of the perforant pathway; the tract connecting the entorhinal cortex with the hippocampal formation as part of the Papez circuit, in late stages of ALS [35, 64]. This is concomitant with the presence (not severity) of pTDP-43 accumulation and neuronal degeneration in the dentate gyrus area in pure ALS patients with memory impairment [59, 60]. Here, regions connected by the perforant path were affected; laminar spongiosis was found in the dentate gyrus and in two severe cases also in the subiculum, as was visualised with standard haematoxylin and eosin staining. The spongiosis in the dentate gyrus may be the result of degenerating axons (originating from the entorhinal cortex) that no longer terminate in the molecular layer of the dentate gyrus. However, white matter alterations were not directly investigated in this study.

**Figure 1.**
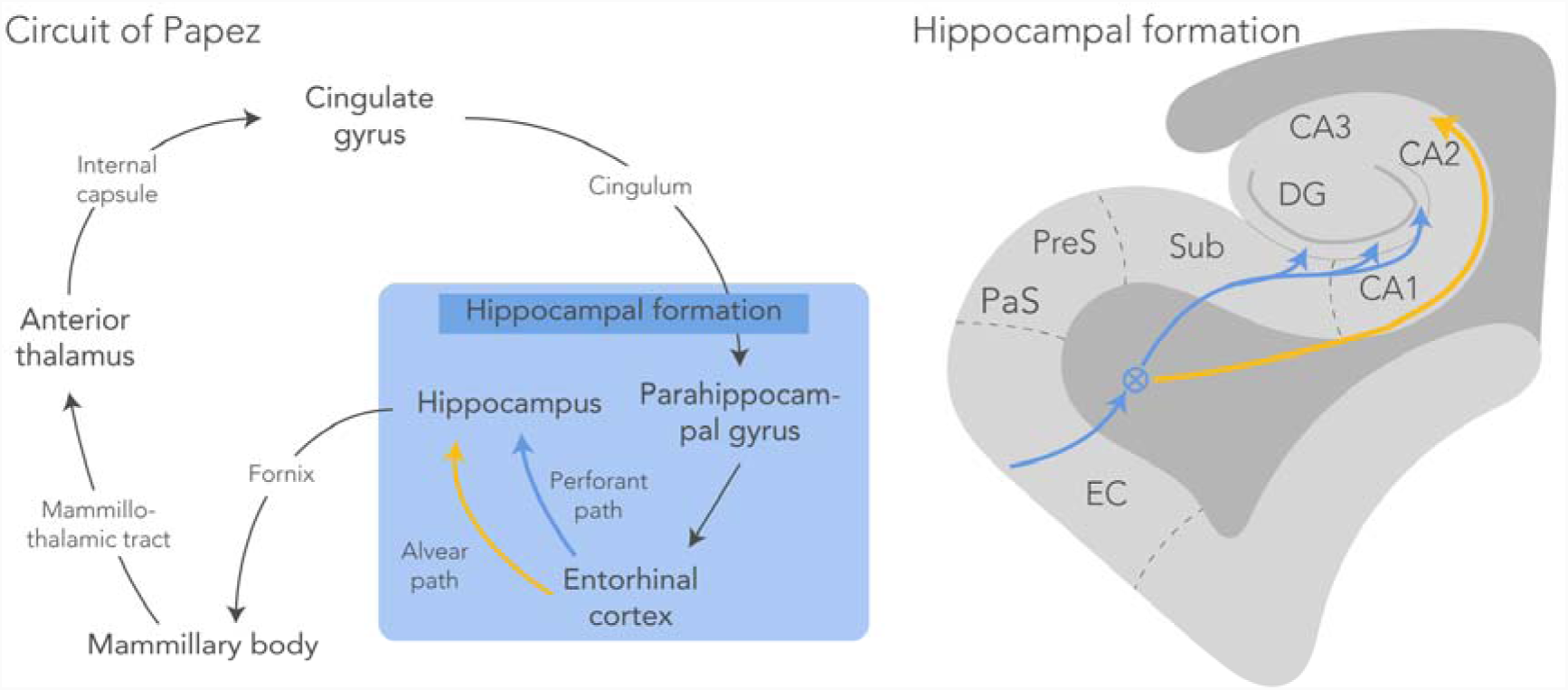
Circuit of Papez with a schematic representation of the hippocampal formation. The perforant path (blue) starts in the entorhinal cortex and projects through the subiculum via the molecular layer of the hippocampus to the dentate gyrus and the CA3 region of the hippocampus (circled-cross indicates out of screen – i.e., anterior posterior – orientation). The alvear path (yellow) projects via the CA1 region into the hippocampus. CA, cornu ammonis; DG, dentate gyrus; EC, entorhinal cortex; PaS, parasubiculum; PreS, presubiculum; Sub, subiculum.

In recent years, advances in diffusion Magnetic Resonance Imaging (dMRI) have attracted researchers to study white matter tissue integrity non-invasively. dMRI infers tissue microstructure from the diffusion profile of water within tissue [11]. The diffusion profile is generally characterised using the metrics mean diffusivity (mean apparent diffusion across all directions), fractional anisotropy (the degree of diffusion directionality), and the principal diffusion direction. These metrics are sensitive to changes in nervous tissue (e.g., demyelination) following neurodegenerative disorders or traumatic brain injury [3]. As such, dMRI is currently a widely accepted imaging modality for studying patients with various neurodegenerative diseases (see Goveas *et al*. for an excellent overview [24]). However, axon density, packing and myelination all alter the diffusion signal [8], muddying the interpretation of dMRI signal changes. To truly identify the origin of dMRI signal changes, histological examination of tissue remains essential. Therefore, ex-vivo dMRI is rapidly evolving to study brain tissue post-mortem and facilitates subsequent comparison to microscopy [49]. A further advantage is the use of high-field MRI systems and longer acquisition times, that offers imaging at much higher resolution with ex-vivo dMRI. With the estimation of fibre orientations at each voxel, dMRI tractography allows one to reconstruct white matter bundles. Application of in-vivo dMRI tractography to delineate the perforant path is challenging because of limited spatial resolution to capture its small size. The use of high-resolution ex-vivo dMRI previously showed to be suitable for reconstructing the perforant path in post-mortem hippocampus specimens [4].

To assess the white matter alterations in the perforant path of patients with ALS, we propose to combine ex-vivo dMRI with subsequent analysis of the microstructural changes using immunohistochemistry (IHC) and polarized light imaging (PLI). PLI quantifies myelin density and the orientation of axons based on birefringence of the myelin sheath [7]. The comparison was made in post-mortem hippocampus specimens from ALS patients and healthy controls. We hypothesize that white matter degeneration of the perforant path – as part of the Papez circuit – in the hippocampus is a feature of ALS, even in the absence of fully developed FTD or deposition of pTDP-43 inclusions in hippocampal granule cells.

These results may provide insight into the earliest neural substrate of FTD-like symptoms observed in ALS and contribute to the broader hypothesis that ALS and FTD may form part of a disease continuum.

## Methods

### Tissue specimens

For this study, fourteen post-mortem ALS human brains were acquired from the Oxford Brain Bank, Nuffield Department of Clinical Neurosciences, University of Oxford, United Kingdom. Five age-matched post-mortem human control brains were acquired via the body donor program at the Department of Anatomy of the Radboud University Medical Center, Nijmegen, Netherlands. The brains were removed from the skull and fixed in 10% formalin. Details on the history of each specimen are listed in Table 1. All brains were cut in coronal slabs of 10 mm thickness and the hippocampi were sampled from a slab comprising the lateral geniculate nucleus as a key anatomical landmark. ALS classification was based on the neuropathological assessment of pTDP-43 proteinopathy in gatekeeper regions previously defined by Tan *et al* [61]. To assess whether the subjects also demonstrated signs of FTD pathology, we checked for pTDP-43 pathology in the dentate gyrus of the hippocampus specimens [61]. Representative blocks of thirteen other brain regions (Supplementary Table 1) were sampled and stained following standard brain banking protocol for diagnostic purposes [42].

**Table 1.**
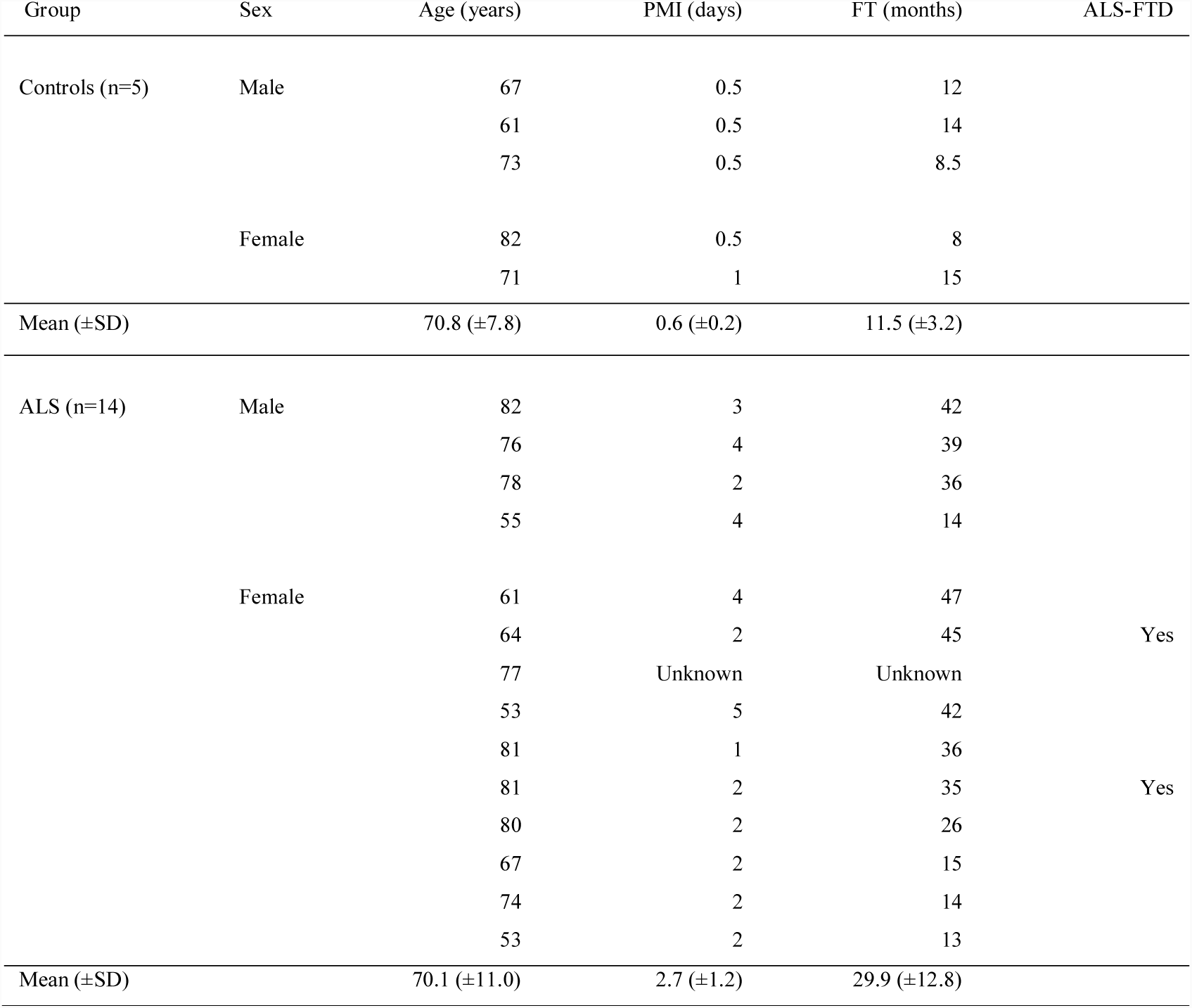
Post mortem specimen details from control and ALS brains. DG, dentate gyrus; FT, fixation time; PMI, post-mortem interval; SD, standard deviation.

### MRI acquisition

Prior to scanning, all specimens were drained of formalin and rehydrated in phosphate buffered saline (PBS) (0.1M, pH=7.4) for at least 24 hours to increase the T2-relaxation rates close to in-vivo levels. Formalin fixation is known to reduce T2-relaxation rates of tissue; thus lowering the MRI signal [53]. Following rehydration, the specimens were placed in a 100 mL syringe filled with fluorinert® (Solvay Solexis Inc), a proton-free liquid which is susceptibility matched to brain tissue. All scans were performed at room temperature on a 11.7T Bruker BioSpec Avance III preclinical MR system (Bruker BioSpin, Ettlingen, Germany) using a birdcage coil (Bruker Biospin). Anatomical scans were acquired with a T1-weighted fast low angle shot gradient echo (FLASH) sequence at a resolution of 0.1×0.1×0.1 mm (TR=25 ms, TE=3.4 ms, flip angle=10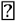).

Diffusion weighted images were obtained using a spin-echo sequence with an EPI readout in four segments (TR=13.75 s, TE=30.1 ms, Δ=12.5 ms, δ=4.0 ms). 64 gradient directions were acquired at b=4000 s/mm^2^, in addition to 6 images with negligible diffusion weighting (b=14 s/mm^2^). The specimens were covered in the coronal plane of the scanner with a field-of-view of 28.8 × 28.8 mm, 72 × 72 matrix, and slice thickness of 0.4 mm, yielding 0.4×0.4×0.4 mm resolution. The number of slices varied between 80 and 100 depending on the size of each specimen.

### MRI analysis

Processing of all MR images was performed with FSL [33]. First, eddy current correction was performed using EDDY and each volume was aligned using a 3D affine transformation. A diffusion tensor was then estimated at each voxel in the dMRI data. Four different microstructural metrics were derived from this diffusion tensor fractional anisotropy (FA), mean diffusivity (MD), axial diffusivity (AD) and radial diffusivity (RD). These metrics are broadly known to be sensitive to tissue microstructure, but not to be highly specific to a single microstructural property. FA describes the diffusion directionality of water ranging from 0 (no directionality of water diffusion, i.e. isotropic diffusion) to 1 (water diffusion along a single direction only). FA in white matter is generally high, but may be decreased in the presence of axonal/myelin degeneration. MD is a measure for the average diffusivity in a voxel and is inversely sensitive to necrosis, oedema and membrane permeability [12]. AD and RD disentangle the FA into diffusivity parallel and perpendicular to the fibre orientation, respectively. An increase in RD was previously shown to correlate with a reduction in myelin [5], but would also be expected to increase if fibre density decreases.

To delineate the perforant path in the hippocampus specimens, we performed probabilistic tractography using the dMRI data, after which we extracted its average microstructural metrics. First, we estimated the fibre configuration in each voxel using the Bedpostx algorithm [10]. We fitted up to three fibre orientations within each voxel in a data-driven way that only estimates additional fibres if their existence is statistically supported by the data. Probabilistic tractography was then performed using the probtrackx2 algorithm [9]. The subiculum and the dentate gyrus were manually segmented and served as seed and termination masks, respectively. The part of the perforant path that runs from the entorhinal cortex to the subiculum (Fig. 1) is located more posterior and was not included in our specimens. 5000 streamlines were generated for each seed-voxel, and only streamlines that entered the dentate gyrus were saved. The curvature threshold was set to 0.2, with a step length of 0.1 mm and a minimal fibre volume threshold of 0.01. A loop check was performed such that pathways that looped back to themselves were excluded. The approach above produced probability maps of the spatial trajectory of the perforant path. An empirically defined threshold was set for the probability maps of each hippocampus separately to obtain a mask of the perforant path. These thresholded maps serve as regions-of-interest (ROI) from which the microstructural metrics were extracted.

Finally, partial volume effects (i.e., the mixture of multiple tissue type within a voxel) can have a significant impact on the diffusion signal [1]. To measure partial volume effects in each specimen, two tissue types (i.e., grey and white matter) were segmented using FSL’s FAST algorithm. FAST then provided partial volume estimates for the tissue types in each voxel. Using the ROIs defined by tractography, partial voluming was extracted from the perforant path.

### Tissue processing

After dMRI, all nineteen hippocampus blocks were bisected along the coronal plane into two approximately equal parts. One part was processed for paraffin embedding for IHC analysis. The other part was frozen for PLI after cryoprotection with 30% sucrose in PBS. Frozen sections are preferred for PLI as the lipids in the myelin sheath remain preserved.

### Polarized light imaging

Using polarized light imaging (PLI), we can estimate the orientation and density of myelinated fibres in thin tissue sections, due to the interaction of polarized light with myelin birefringence. The theory behind the polarimetric set-up is extensively described elsewhere [7, 38].

All frozen tissue blocks were sectioned at 50 µm thickness on a freezing microtome (Microm HM 440E Microtome). The sections representing the perforant path were mounted on glass slides and coverslipped with polyvinylpyrrolidone, a water-soluble mounting medium. PLI images were acquired on a Zeiss Axio Imager A2 microscope upgraded with a stationary polarizer, a quarter wave plate and a rotating polarizer. Light first passes through the stationary polarizer and a quarter wave plate positioned in a 45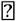 angle relative to the stationary polarizer to create circularly polarized light. The polarized light then passes through the hippocampal sections undergoing a phase shift caused by birefringence of myelin, and finally through the rotating polarizer. To quantify the phase shift, which reflects the in-plane orientation of fibres, images were acquired at 9 equiangular orientations of the rotating polarizer from 0° to 160° with a CCD camera (Axiocam, Zeiss). Together with a 1.0x magnifying objective, this yielded a spatial resolution of ∼4 μm/pixel. A set of background images was acquired for every rotation angle to correct for inhomogeneous background illumination [20]. Three parameter maps were derived by fitting the light intensity at each pixel to a sinusoidal function [34]. The phase of the sinusoid gave the in-plane orientation of myelinated fibres. The amplitude of the light wave provided the retardance map describing the myelin density. The retardance and in-plane orientation maps are combined in the fibre orientation map that colour codes the direction fibres with its intensity modulated by the myelin density (Axer *et al*., 2011b). Finally, a transmittance map was calculated as the average amount of light passing through the tissue yielding grey and white matter contrast.

The retardance maps inform about how much the phase of the light wave is shifted due to interaction with myelin and is therefore used to evaluate the amount of myelin within a section. A decrease in the retardance values may therefore be a marker of myelin loss. The perforant path was manually delineated using a composite of the PLI transmittance and retardance maps for each specimen separately. This was defined as the white matter originating from the subiculum and partly projecting through CA1, along and into the dentate gyrus (Fig. 3).

### Immunohistochemistry

IHC staining of the hippocampus sections was carried out to identify disease specific changes in microstructure. Antibodies against several proteins were used; proteolipid protein (PLP) for myelin, SMI-312 for axonal phosphorylated neurofilaments, cluster of differentiation 68 (CD68) for activated microglia and lastly pTDP-43. Paraffin embedded tissue blocks were sectioned at 6 µm for the PLP, SMI-312 and CD68 stains. Sections stained for pTDP-43 were cut at 10 μm. The sections were then mounted on Superfrost Plus slides (ThermoFisher, Art. No. J1800BMNZ). The sections were deparaffinised in xylene and rehydrated through a series of 100%, 100%, 90% and 70% ethanol. The endogenous peroxidase was blocked by incubating the sections 3% H_2_O_2_ in PBS for 30 minutes. Subsequently, antigen binding sides for pTDP-43 and CD68 were retrieved by autoclaving in citrate buffer (pH=6). Antigen retrieval for PLP and SMI-312 was achieved through heating the sections submerged in citrate buffer (pH=6) in a microwave. After antigen retrieval the sections were washed with PBS. After washing, the primary antibody was added. All primary antibodies were diluted in TBS/T (pH=7.6). The primary antibodies against PLP (1:1000, Bio-Rad, MCA839G), CD68 (1:200, Dako Denmark, M0876) and SMI-312 (1:2000, BioLegend, 837901) were incubated for 1 hour at room temperature. The primary antibody against pTDP-43 (1:40 000, CosmoBio, TIP-PTD-M01) was incubated over night at 4°C. After washing, the secondary antibody containing HRP rabbit/mouse serum (DAKO EnVision Detection Systems) was added and incubated for 1 hour. A DAB substrate buffer solution was subsequently added (DAKO EnVision Detection Systems) and incubated for 5 minutes. Sections were washed with PBS to stop the reaction. Finally, a counterstain for haematoxylin was performed to visualise the nuclei. After counterstaining, the sections were washed, dehydrated and mounted with DPX.

IHC sections were digitized on an Aperio ScanScope AT Turbo slidescanner employing a 20x magnifying objective (40x for the pTDP-43 sections). pTDP-43 sections were judged by two neuropathologists (OA and MPG) to determine any TDP-43 pathology in the dentate gyrus for FTD-TDP proteinopathy. The other stains (PLP, SMI-312 and CD68) were computationally segmented in Matlab (MathWorks Inc., Natick, USA) making use of colour contrast between the DAB stain and the haematoxylin counterstain. Following common practice in IHC analysis, the area fraction of the IHC stains was used to quantify the protein of interest. To achieve this, the images were first transformed to HSV colour-space. A colour range for the brown DAB staining was interactively determined in the hue-channel (H) using Matlab’s colour-segmentation application. Furthermore, an intensity threshold was applied in the saturation-channel (S). The colour and intensity threshold were used together to isolate the brown DAB-staining from the background. However, to account for varying staining intensity across specimens, the intensity thresholds were determined for each specimen separately. The intensity threshold was automatically determined by measuring the average intensity in the saturation-channel from a region within the collateral white matter in each specimen. Non-tissue pixels were likewise identified as bright white pixels that were not labelled DAB-staining, nor the haematoxylin counterstain. Finally, a stained area fraction was calculated as the ratio of DAB positive pixels over all tissue-pixels within a local neighbourhood of 100×100 pixels. The perforant path area was manually drawn in all slices separately. Following a similar strategy as with PLI, these regions-of-interest were defined as the white matter originating from the subiculum and partly projecting through CA1, along and into the dentate gyrus.

### Statistics

All statistical analyses were carried out in Matlab. We constructed a general linear model (GLM) that tested for group differences (similar to a student’s t-test) among the dMRI microstructural metrics (FA, MD, RD and AD). To assess whether microstructural differences were driven by partial volume effects, the GLM included the tissue partial voluming estimates from the FAST algorithm as a covariate. Between the groups there was a significant difference in fixation time (p=0.003). Since it is known that fixation time and post-mortem interval can cause a reduction in FA and diffusivity [19, 57], a normalization step was performed before calculating the t-statistics. The normalisation was done by dividing the found dMRI metrics in the perforant path by a reference value for each of these metrics in a control region – collateral white matter – within the same sample.

A one-tailed student’s t-test was applied to evaluate PLI retardance values in the perforant path. To assess whether changes in dMRI outcomes were driven by myelination, a Pearson correlation (two-tailed) was performed between both FA and MD, and PLI retardance. Finally, one-tailed t-test was again used quantify differences in IHC staining between ALS and controls. PLP myelin estimates were correlated against SMI-312 neurofilaments, PLI retardance and dMRI FA (Pearson, two-tailed).

With ALS and FTD forming a disease spectrum, the heterogeneity of metrics within the disease group is expected to be large. As such, the correlation analyses may unravel relationships that extend across all subjects; for example, between dMRI and myelin integrity.

## Results

### MRI

Diffusion MRI was acquired from all subjects, but one ALS case was excluded from further analysis due to insufficient contrast and noisy images. The perforant pathway was reconstructed from the dMRI data by employing probabilistic tractography. The probability maps of the perforant path from two control and two ALS cases are depicted in Figure 2A. Tracts are shown from the subiculum to the dentate gyrus; the part from entorhinal cortex to subiculum was not included in the hippocampus specimens. An arbitrary probability threshold was applied for specimen separately to create the maps. These regions defined by the probability maps were used to extract the diffusion tensor metrics and to get an impression of white matter degeneration along the perforant paths in ALS cases and controls (see Fig. 2B).

**Figure 2.**
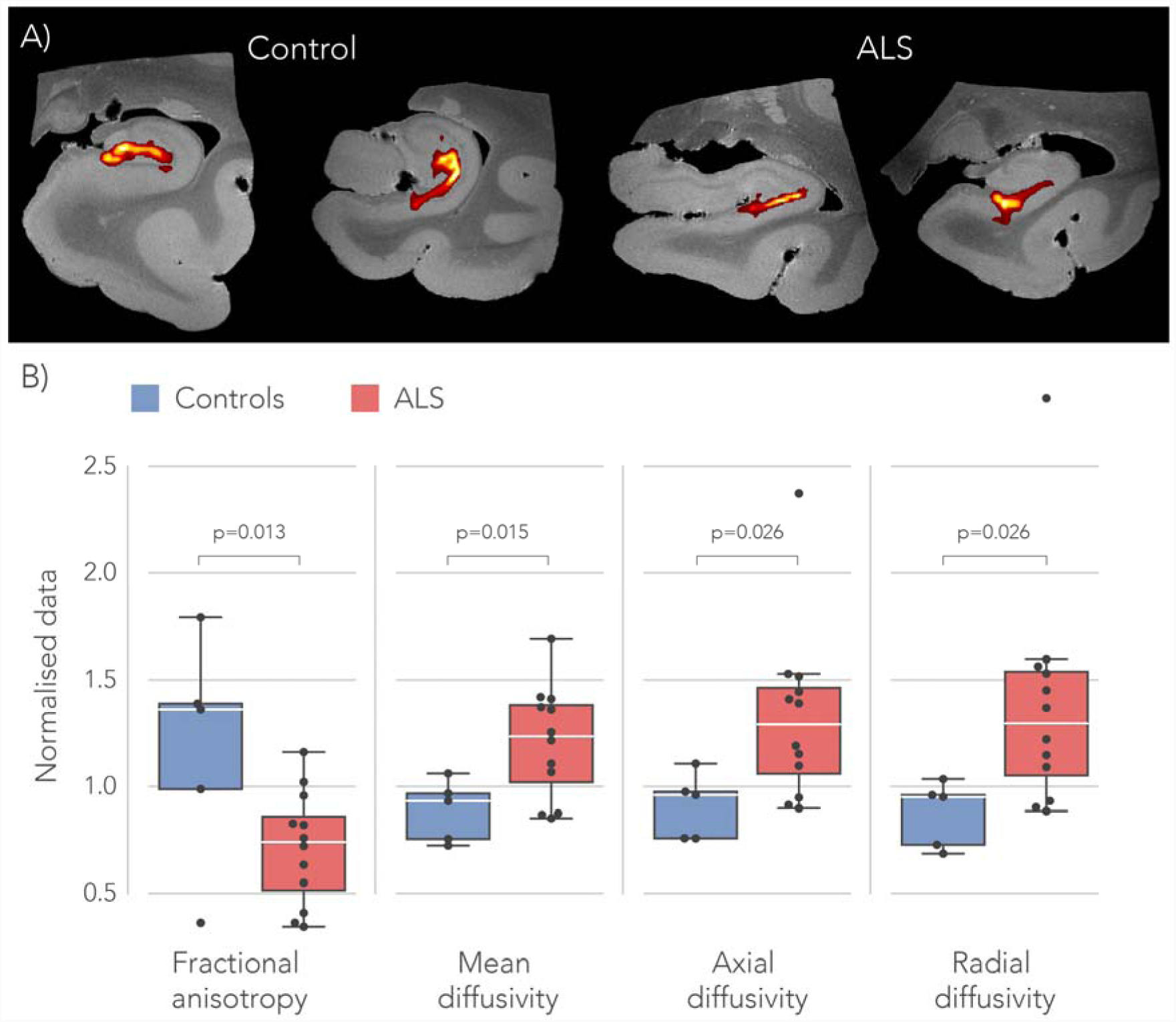
Perforant path microstructure estimated from dMRI data. **A)** Tractography in the perforant pathway running from the subiculum to the dentate gyrus for two controls and two ALS cases. These tracts were used for extracting the MD, RD, AD and FA values within the perforant path. Tracts are superimposed on the T1-weighted anatomical scan. **B)** The FA value in the part of the perforant path that runs from the subiculum to the dentate gyrus is lower in ALS cases compared to controls (p=0.013). The MD value is significantly higher in the perforant path in the ALS specimen compared to controls (p=0.015) as well as the AD (p=0.026) and the RD (p=0.026). All p-values given for a one-tailed t-test. The boxplot centre lines depicts the median for each metric; box limits, the 25th and 75th percentiles of the metrics; the whiskers extend to the most extreme data points excluding outliers; raw data points are given by the black dots.

Within the perforant path, the FA values were significantly lower in ALS specimens compared to controls (p=0.013). A reduced FA is caused by water diffusing more isotropic (less directional) and possibly indicates myelin degeneration or an increase in interstitial spaces. This hypothesis is further strengthened by the increase that was found in MD in ALS specimens compared to controls (p=0.015). In addition, both AD (p=0.026) and RD (p=0.026) values were significantly higher in ALS cases compared to controls. An increase in RD might be caused by an increasing amount of diffusion along the radial diffusion direction, again consistent with loss or atrophy of fibres. The model including partial volume effects did not identify a significant effect of partial volume (0.07<p<0.45); moreover, inclusion of partial volume as a confound regressor did not change the pattern of a significant difference between groups reported in Fig. 2.

### Polarized light imaging

Figure 3 depicts an example of the transmittance, retardance and fibre orientation maps from a control sample. White matter is easily recognized in the retardance maps, appearing bright due to birefringence in the myelin sheath. Several sub-areas of the hippocampal formation are delineated in the fibre orientation map, including the approximate trajectory of the perforant path.

**Figure 3.**
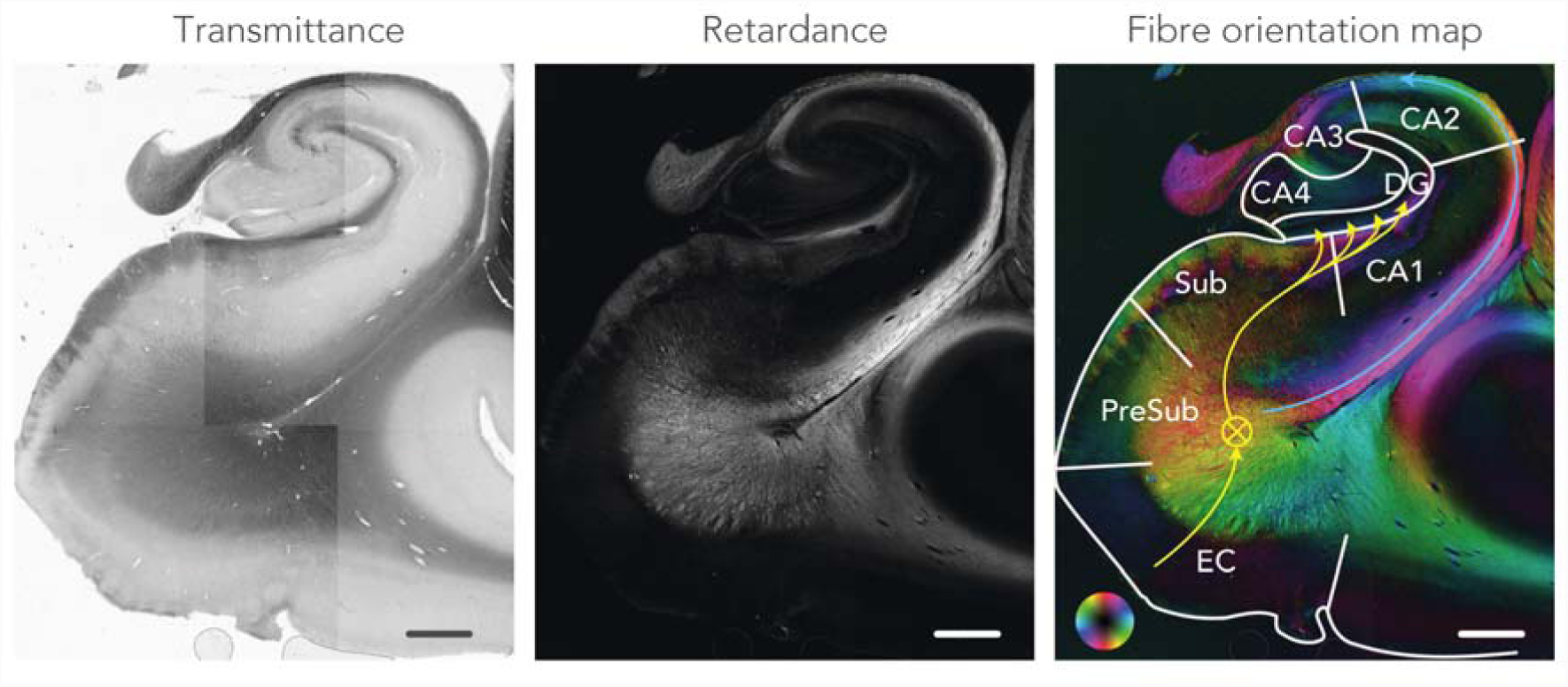
Polarized light imaging data from a control case. The transmittance map reflects the amount of light passing through the tissue. The retardance map shows the degree to which the light is shifted due to interaction with the myelin. The fibre orientation map colour-codes the directionality of the myelinated fibres indicated by the colour wheel. This map is formed by combining the retardance and in-plane map. The anatomical regions within the hippocampus are delineated and the approximate direction of the perforant path is indicated with the yellow arrows. Circled cross represents anterior-posterior direction of the perforant path. The maps are a composite of multiple field-of-views. CA, cornu ammonis; DG, dentate gyrus; EC, entorhinal cortex; PreSub, pre-subiculum and Sub, subiculum. Scale-bar = 3 mm.

PLI retardance maps were used as a semi-quantitative measure of myelin density. Example retardance maps from a control case and an ALS case are shown in Figure 4, demonstrating lower retardance values in the perforant path area for the ALS case. Across all samples the retardance values appeared to be lower in the perforant paths in ALS (p=0.05). The correlation analysis revealed a significant association between FA and retardance (r=0.52, p=0.04). This may indicate that reduced FA is driven by demyelination of the perforant path. No significant correlation was found between retardance and MD (r=0.34, p=0.48).

**Figure 4.**
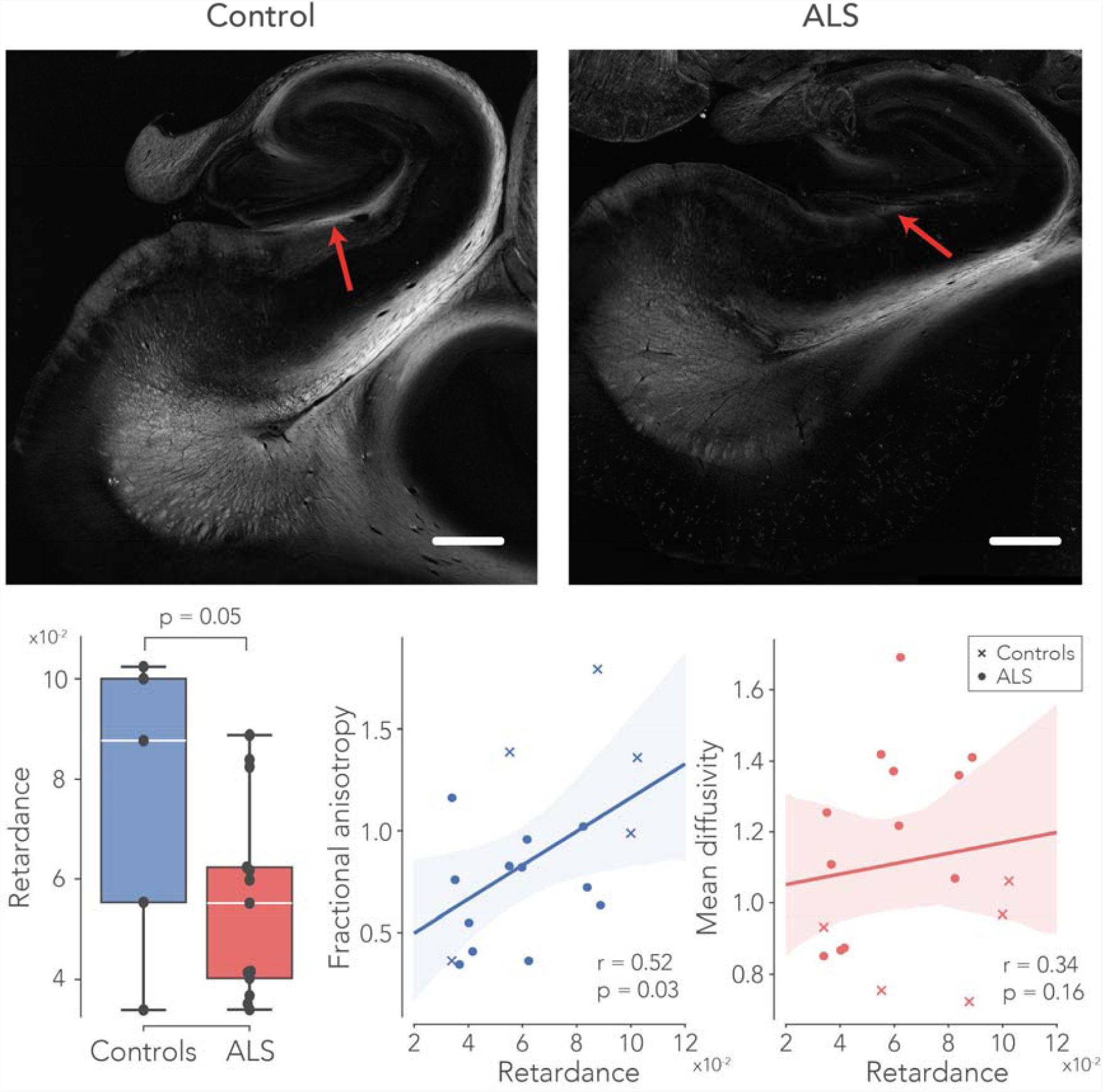
PLI retardance – reflecting myelin density – in the perforant path. The upper part of the figure depicts two exemplar retardance maps from a control – and an ALS case. Here, the red arrows indicate the approximate location of the perforant path. Retardance was significantly lower in ALS patients compared to controls. Furthermore, the changes in FA were possibly driven by myelin density, supported by significant correlation between FA and retardance were significantly correlated across all subjects. A correlation between MD and retardance was not found. See caption Figure 2 for boxplot description. Scale-bar = 3 mm.

### Immunohistochemistry

IHC was successfully carried out in all hippocampus specimens. Two out of the fourteen ALS cases demonstrated FTD-TDP proteinopathy in the dentate gyrus (Fig. 5). The automatic segmentation of the myelin (PLP), neurofilaments (SMI-312) and activated microglia (CD68) images was deemed suitable for semi-quantification of these IHC markers (Fig. 6). Following up on the white matter alterations reported with dMRI and PLI, levels of myelin and neurofilaments were compared between controls and ALS patients. Indeed, less myelin was found in the perforant path of ALS patients (p=0.03). Also, the stained area fraction of neurofilaments was lower in ALS (p=0.02). No changes were observed in the amount of CD68 in both groups (Fig. 7). To see if reduced levels of PLI retardance and dMRI FA were driven by lowered levels of myelin, a correlation analysis across all specimens was performed. Neurofilaments area fractions were lower in subjects with reduced myelin area fraction (r=0.71, p=0.001). Additionally, a positive correlation was found between IHC myelin and (1): PLI retardance (r=0.51, p=0.03) and (2): dMRI FA (r=0.52, p=0.03). The two ALS-FTD cases expressed lowered levels of myelin in comparison to all other ALS cases, but these were not significantly different (Fig. 7). CD68 area fractions in ALS-FTD were also similar compared to other ALS cases.

**Figure 5.**
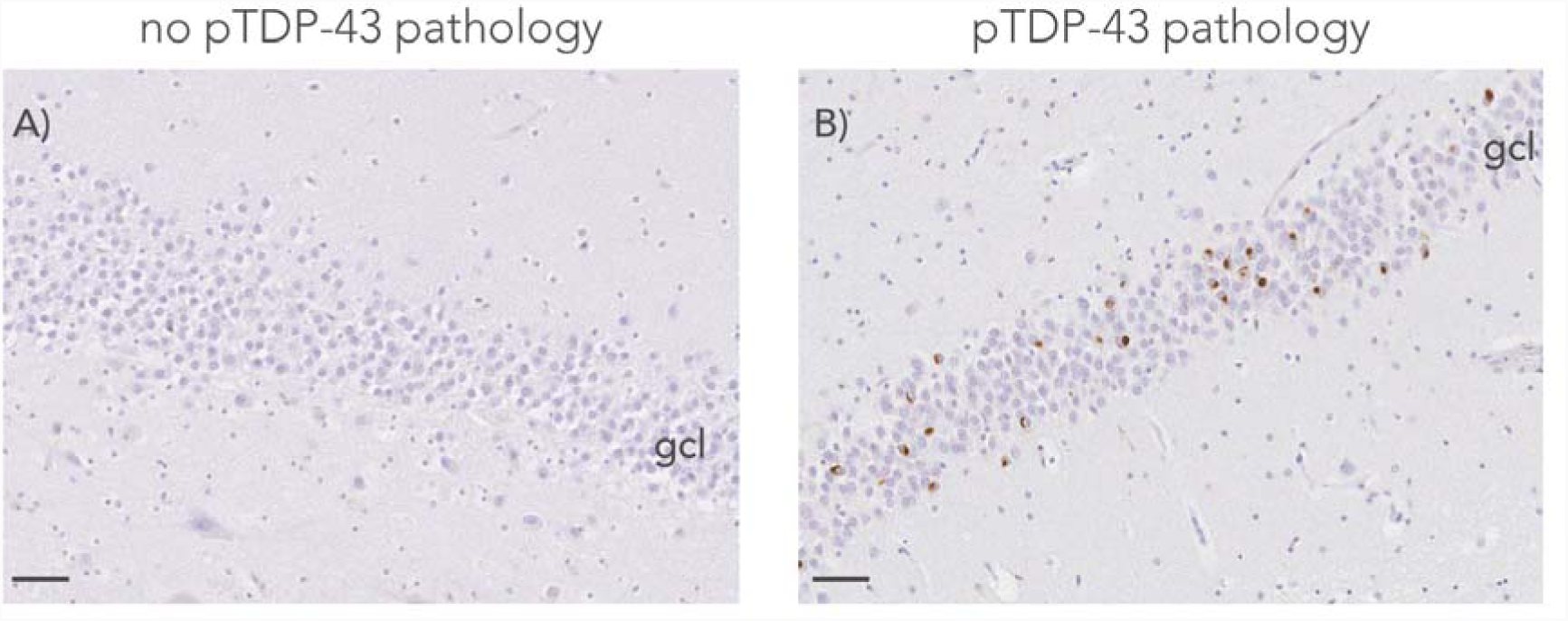
pTDP-43 pathology in the granule cell layer (gcl) of the dentate gyrus. Cases were stratified by absence **(A)** or presence **(B)** of pTDP-43 inclusions (coloured brown) in the granule cells of the dentate gyrus. Blue nuclei. Scale-bar = 50 μm.

**Figure 6.**
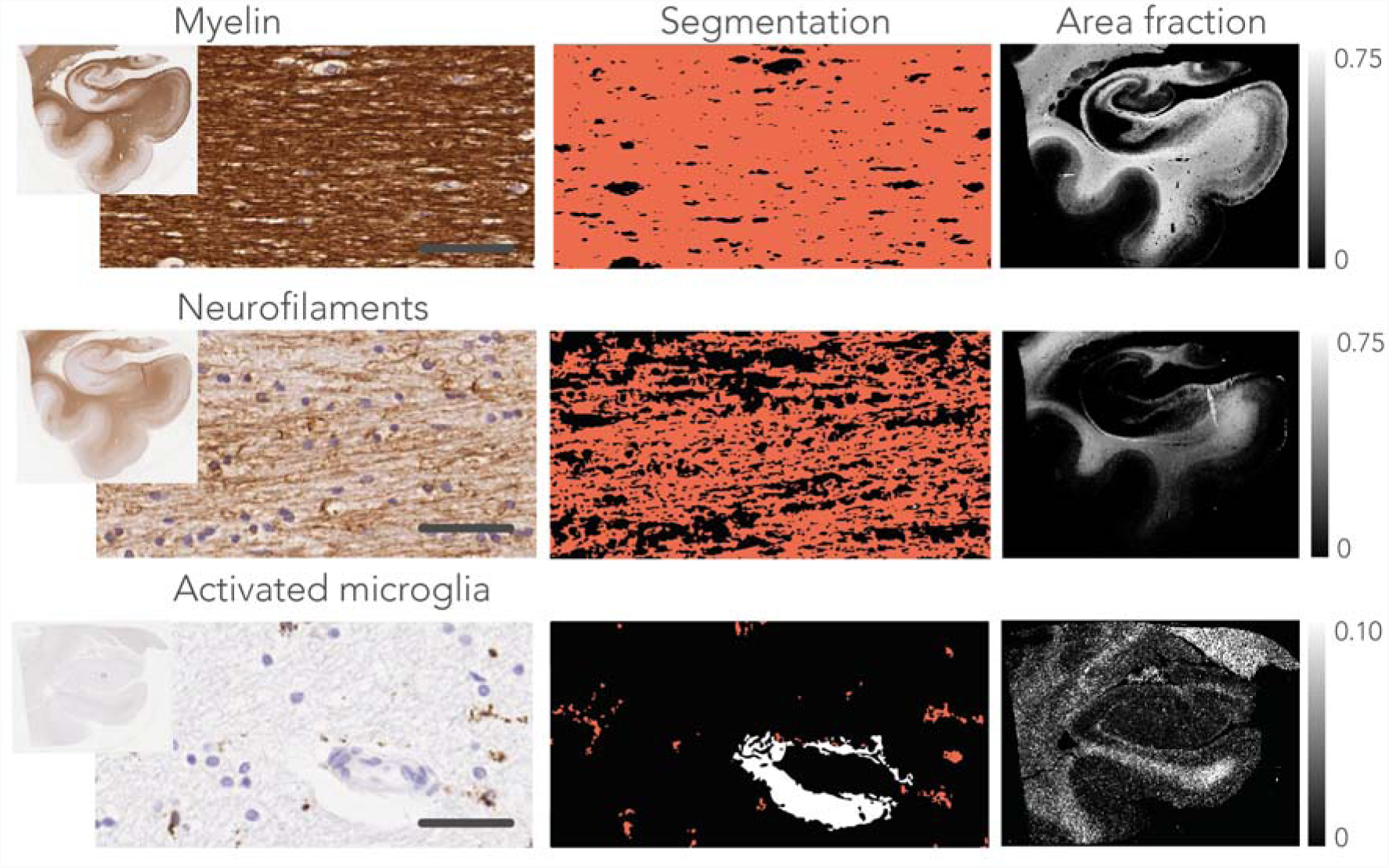
Histological data processing. The left column shows stained sections with a snapshot for myelin (PLP), neurofilaments (SMI-312), and activated microglia (CD68). The middle column depicts the segmentation of the images in the left column. A colour-based segmentation was applied to separate DAB-stain positive pixels (red) from the background tissue (black) and non-tissue pixels (white). After processing, the stained area fraction was calculated per 100×100 pixels resulting in a down-sampled image reflecting the stained area fraction over the entire slice (right column). Scale-bar = 30 μm.

**Figure 7.**
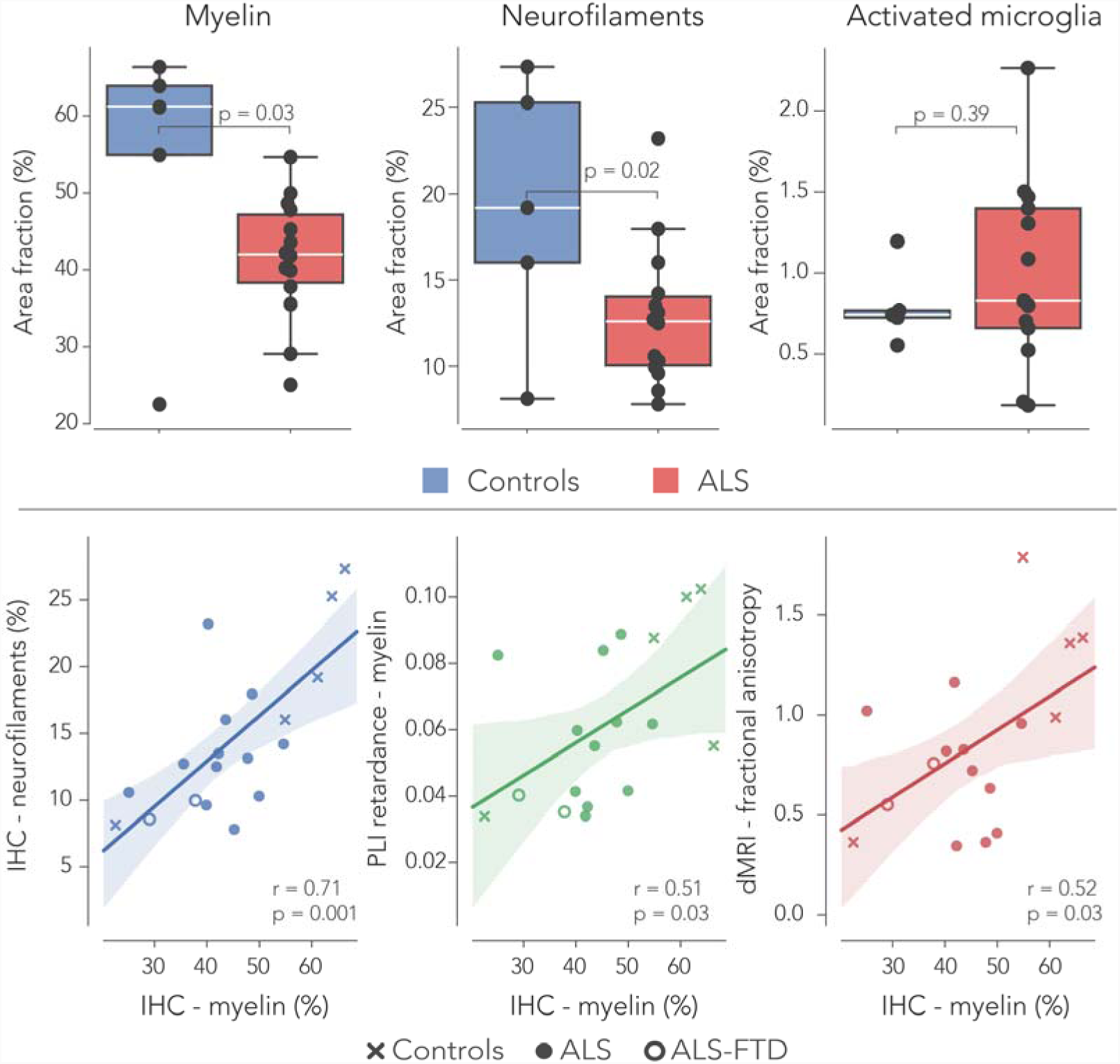
Immunohistochemistry in ALS and controls. The stained area fraction of myelin (PLP) and neurofilaments (SMI-312) was significantly lower in the perforant paths of ALS patients compared to controls. The area fractions of activated microglia (visualised with CD68; marker for inflammation) was similar in both groups. Two out of the fourteen ALS cases exhibited pTDP-43 pathology within the dentate gyrus, used for neuropathological characterisation of FTD. Furthermore, myelin area fractions from the IHC stains correlated with myelin density estimated from the PLI retardance maps and also with FA derived from dMRI. The two ALS-FTD cases had lowered myelin density estimates, but no difference was observed compared to pure ALS cases. See caption Figure 2 for boxplot description.

## Discussion

ALS is mostly known as a motor neuron disease; the corticospinal tract and the corpus callosum are frequent targets for studies investigating white matter alterations in ALS [17, 22, 31, 40]. However, ALS is increasingly recognized as a multisystem neurodegenerative disorder, that may be associated with clinical bvFTD. As such, white matter alterations were studied in the perforant path, a component of the Papez circuit, as evidence for ALS being multi-system disorder [48].

Using diffusion MRI, we demonstrated a reduction in fractional anisotropy (FA) and an increase in diffusivity markers (MD, AD and RD) in the perforant path in ALS (Fig. 2), all suggesting white matter degeneration (Bozzali *et al*., 2002; Soares *et al*., 2013). However, in the presence of demyelination and white matter loss, AD has also been reported to remain unaffected (Song *et al*., 2002; Klawiter *et al*., 2011) or even decrease (Tyszka *et al*., 2006; Harsan *et al*., 2006). Possibly, an increase in AD is observed due to enlargement of the extracellular space as a result of axonal degeneration. While the dMRI results suggest that white matter in the perforant path is damaged in ALS cases, changes in the dMRI metrics can also be driven by several other factors. dMRI microstructure estimates are inferred from local diffusion of water in tissue and are therefore indirect measures of the white matter geometry. Rather than an altered microstructural signature due to disease, it is possible that some changes are driven by variations in anatomy across subjects. Furthermore, death, tissue fixation and post-mortem interval all affect the microstructural metrics estimated here. For instance, a significant difference was observed in fixation times between the control and ALS group (p=0.003), which is known to decrease diffusivity in tissue [19, 50, 51, 57]. We accounted for these fixation time differences by normalizing the dMRI measures with a control region in collateral white matter.

Partial volume effects (PVE) – reflecting intravoxel heterogeneity of tissue types – can also be a significant confounding factor in age or disease related dMRI analyses [63]. For ex-vivo dMRI we expected this to be less an issue, because the spatial resolution is high (0.4 mm isotropic). Indeed, after estimating PVE in the perforant path and including it in our modelling to evaluate microstructural group differences, PVE had no significant effect on the outcomes (Fig. 2).

Finally, as dMRI measures are affected by death and fixation, one ideally would perform in-vivo dMRI of the perforant path in control and disease groups. Using current clinical MR systems – typically having field strengths of 1.5-3T – this remains challenging; with limited scan times (< 1 hour) for patients, obtaining sufficient resolution to image small structures such as the perforant path is currently infeasible with dMRI. The use of ex-vivo tissue allowed us to employ long scan times (3-4 hours) and an ultra-high field MRI scanner (11.7T) resulting in very high resolution dMRI. Nevertheless, promising progress in this area is currently made by obtaining in-vivo dMRI with whole-brain coverage at sub-millimetre resolution [52].

Polarized light imaging is a relatively novel imaging modality and to date primarily applied to study neuroanatomy [6, 28], interpret dMRI fibre orientations in grey matter [39] or evaluate dMRI derived fibre dispersion [41]. In particular, hippocampal fibre systems have been visualised with PLI previously [65], demonstrating excellent use of the technique to study connectivity at mesoscopic scale. However, its application to study white matter degeneration is, to our knowledge, new and its results should therefore be interpreted with care. We found a decrease in retardance values in the perforant paths of ALS specimens (Fig. 4). Retardance reflects myelination and thus a reduction could indicate demyelination. However, PLI retardance is not exclusively sensitive to birefringence of the myelin sheath; it also decreases with an increasing fibre inclination (out-of-section) angle. Theoretically, this poses the opportunity to reconstruct the 3D fibre orientation at microscopic resolution, an approach that was adopted by 3D-PLI [7]. 3D-PLI uses a tilting specimen stage that can separate fibre inclination from birefringence. The PLI microscopy set-up employed in this study is not equipped with such a tilting stage and therefore cannot disentangle between variations in fibre inclination and birefringence. Yet, the hippocampal specimens in this study were carefully sectioned in the same orientation keeping variations in fibre inclination angles in the perforant to a minimum. Therefore, we can ascribe differences in retardance primarily to birefringence and thus myelination.

While the dMRI and PLI outcomes suggested white matter degeneration in the perforant path of the ALS specimens, the gold standard remains IHC in these types of analyses. The stained area fractions of both myelin and the neurofilaments were lower in ALS compared to controls (Fig. 7). PLP is one of the most abundant myelin proteins in the central nervous system [14] and demyelination would result in a reduction of PLP estimates [23]. Loss of neurofilaments indicates neuroaxonal damage and its breakdown product in the blood or cerebrospinal fluid is a suitable biomarker for various acute and chronic neurological disorders [36]. A loss of neurofilaments was correlated with reduced myelin density in the perforant path. Also, it is likely that the changes observed in dMRI (lowered FA) and PLI (reduced retardance) are driven by the white matter degeneration of the perforant path; a positive correlation was found between IHC myelin and both dMRI metrics (Fig. 7).

In the hippocampus, or in the granule cell layer of the dentate gyrus in particular, pTDP-43 pathology is present in late stage of ALS and early stages of FTD [61]. Although the two ALS-FTD cases were on the lower spectrum of myelination levels within the ALS group, no evidence was found for extreme white matter degeneration as a result of pTDP-43 pathology in the dentate gyrus. Nevertheless, the fact that white matter degeneration was present in almost all ALS, may suggest that loss of myelin or neurofilaments is present before the onset of pTDP43 pathology in the hippocampus or entorhinal cortex.

IHC analysis showed no differences in the amount of activated microglia – a measure for neuroinflammation – in ALS cases compared to controls. This is somewhat surprising as white matter degeneration is associated with increased microglial activation [44].

Indeed, motor connectivity in the corpus callosum and the corticospinal tract previously showed enhanced levels of microglial activation in ALS (Sugiyama *et al*., 2013; Henkel *et al*., 2004). However, inflammation is a complex system in the brain and there are variety of processes that may drive neuroinflammatory responses [21]. A marker such as IBA-1 (ionized calcium-binding adapter molecule 1) is also frequently applied to estimate neuroinflammation in tissue and may have produced different outcomes [26].

Thus far, only a segment of the perforant path as being part of the Papez circuit was investigated. Future research should also include imaging of the other Papez circuit regions to get a better understanding of the disease’s progression in areas such as the anterior cingulate cortex, fornix, mammillary bodies and the anterior thalamus. Our data would benefit from the inclusion of quantitative cognitive assessment scores for each patient. However, we postulate that neuropathological staging of pDTP-43 pathology can be used as a surrogate marker for the likelihood of progression towards FTD for each case. The current study supports evidence that the perforant path undergoes degeneration, a feature that substantiates evidence for the cognitive deficits observed in some ALS patients. Some more insight might be gained from the relationship between perforant path degeneration and clinically observed cognitive decline.

## Conclusion

This work investigated white matter changes in the perforant path in ALS patients using dMRI, PLI and IHC analyses as a possible marker for FTD development. A reduction in the FA and an increase in the MD, AD and RD observed in all ALS cases relative to controls, implicating white matter degeneration of the perforant path. This observation was supported by a decrease in the PLI retardance values measuring myelin density in the perforant path. IHC confirmed the results from dMRI and PLI by reporting increases in myelin and neurofilaments. We conclude that degeneration of the perforant path occurs in ALS patients and that this may occur before, or independent of, pTDP-43 aggregation in the dentate gyrus of the hippocampus. This provides further evidence of ALS being a multisystem disease, and involvement of the perforant path supports a link between ALS and FTD. Finally, our results warrant future investigations of the perforant path as a candidate neural MRI correlate of the cognitive symptoms in ALS.

## Acknowledgements

We would like to acknowledge “Stichting Alzheimer Nederland” for their funding support. Furthermore, we would like to acknowledge P.J.W.C. Dederen for instructing in the cutting of the sections for PLI. Also, thanks to C. Grabitz and J. de Ruyter van Steveninck for their lab work. Finally, we acknowledge the Oxford Brain Bank, supported by the Medical Research Council (MRC), Brains for Dementia Research (BDR) (Alzheimer Society and Alzheimer Research UK), Autistica UK and the NIHR Oxford Biomedical Research Centre as well as the body Donor program from the department of Anatomy, Radboud UMC Nijmegen for providing the post-mortem brain specimens.

## Data availability

All data is available upon reasonable request.

